## Supplemental Figure 1 for "Neural activity ramps in frontal cortex signal extended motivation during learning"

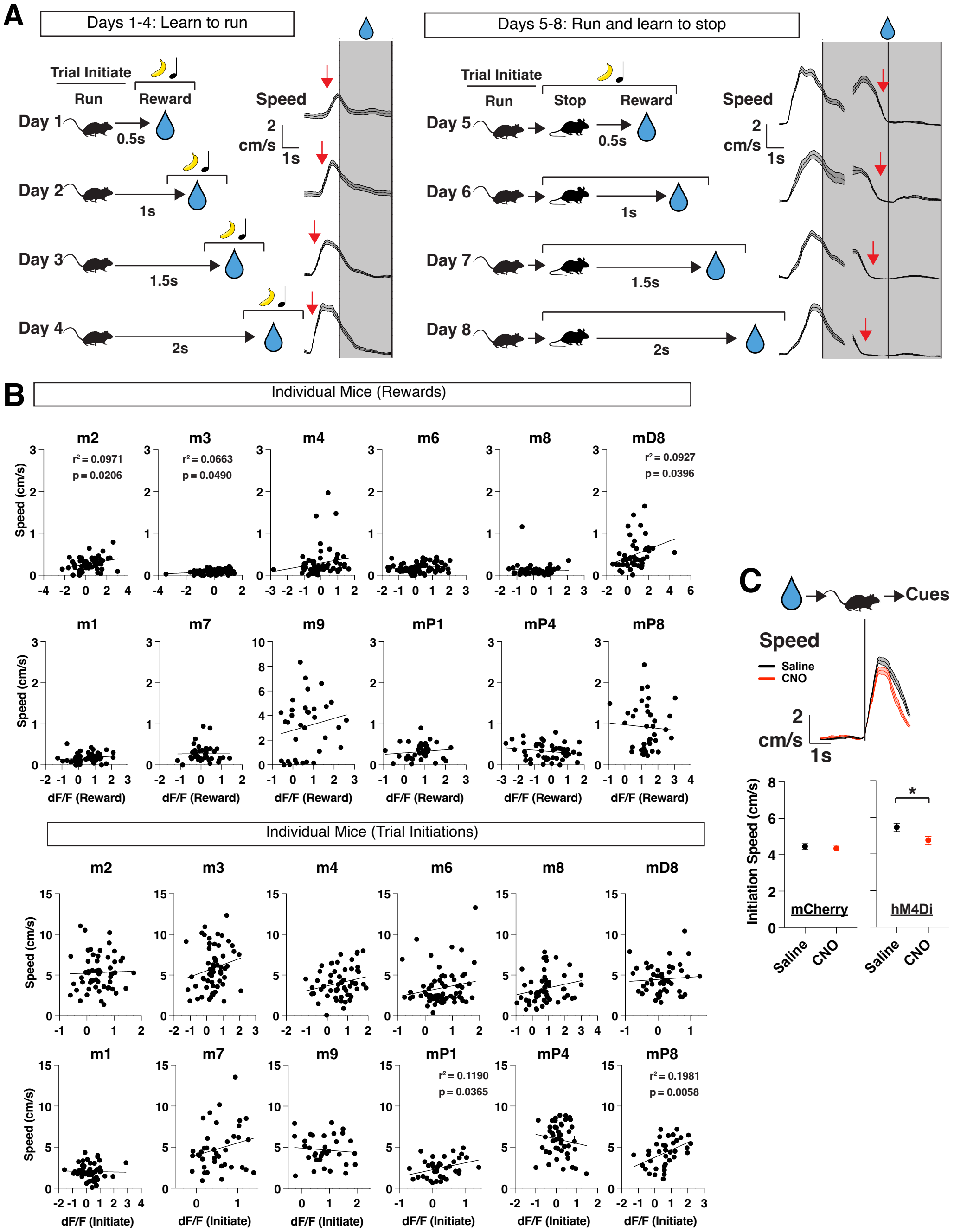


**Figure S1. Task shaping and speed related differences between mice and during ACC inhibition**

(A) Left: schematic of behavioral shaping. Mice were shaped to run (>1 cm/s) for increasing durations over the course of 4 days to obtain rewards. Trial averaged plots of speed along the days of shaping (N=3 mice). Red arrow denotes the increasing duration needed to run to trigger rewards. Right: Days where mice decrease their speed (<1cm/s; stop) during cues for rewards. Red arrows denote the increasing duration to stop to trigger rewards. Data are mean (solid line) ± s.e.m (shaded area).

(B) Scatter plots of speed (cm/s) and ACC dF/F during reward (top) or trial initiations (bottom) for individual mice. Individual data points shown, with a best fit line, represented by the solid line in the figure. r2 and p values, as determined by linear regression, are shown for each mouse that had a p<0.5.

(C) Left: Trial average plots of speed (cm/s) aligned to trial initiation for saline-administered day (black) or CNO (red; N=4 mice). Data are mean (solid line) ± s.e.m (shaded area). Quantification of speed during trial initiation for mCherry-control mice (N=187, 214 trials across 6 mice) and hM4D(Gi)-DREADDs mice (N=166, 120 trials across 4 mice). p=0.5692 for mCherry and *p=0.0217 for hM4Di, unpaired t-test between saline and CNO sessions per group. Data are individual points, with mean ± s.e.m.
