## Supplemental Figure 2 for "Neural activity ramps in frontal cortex signal extended motivation during learning"

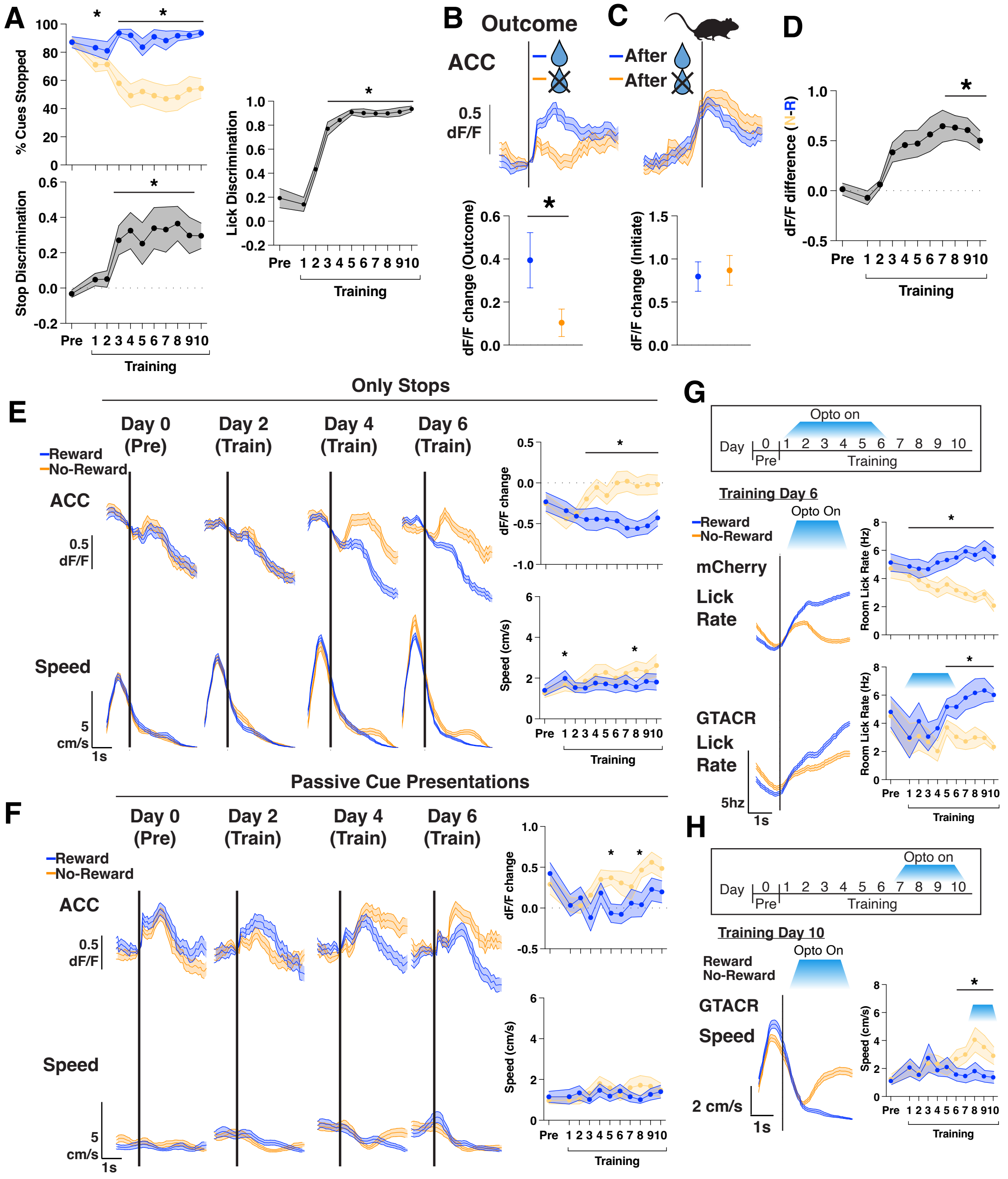

**Figure S2. Lick rate discrimination, ACC learning signal controls, and ACC inhibition lick rate learning**

(A) Left: Percentage of cue presentations with stops, separated by reward and no-reward cues (blue vs orange) and quantification of stop discrimination index (see Methods) across training. Right: Quantification of lick discrimination index (see Methods) across training. N=12 mice, data are mean ± s.e.m. *p<0.05, paired t-test between reward and no-reward each day, or one-way repeated measures ANOVA with post-hoc Tukey's multiple comparison test between each training day and preexposure.

(B) Top: Trial average plots of ACC activity (z-scored dF/F) and speed (cm/s) aligned to outcome onset, separated by reward or no-rewards. Data are mean (solid line) ± s.e.m (shaded area). Bottom: Quantification of the mean dF/F and speed during outcome. N=12 mice in each group, data are mean ± s.e.m. *p<0.05, paired t-test between reward and no-reward each day.

(C) Same as B, but for trial initiations after each outcome.

(D) Quantification of ACC dF/F difference between reward and no-reward cues (see Methods) across training. N=12 mice, data are mean ± s.e.m. *p<0.05, one-way repeated measures ANOVA with post-hoc Tukey's multiple comparison test between each training day and preexposure.

(E) Left: mean dF/F in ACC (top) and speed (bottom) aligned to cue onset for cue presentations in which stops occurred (blue vs orange). N=12 mice. Data are mean (dark line) with s.e.m. (shaded area). Right: Quantification of mean change in dF/F and speed in cue zone, assessed separately for each cue presentation. N=12 mice in each group, data are mean ± s.e.m. *p<0.05, paired t-test between reward and no-reward each day.

(F) Same as E, but for passive presentation of the reward and no-reward cues.

(G) Top: optogenetic inhibition was targeted to days 1-6 of training and mice were allowed to continue training for days 7-10. Bottom left: Trial averaged plots of lick rate (hz) aligned to cue onset on T6 for mCherry controls and GTACR inhibition mice, separated by reward or no reward cues. Bottom right: Quantification of mean lick rate during cues. N=8 mice for mCherry, 4 for GTACR early inhibition. *p<0.05, paired t-test.

(H) Top: optogenetic inhibition was targeted to days 7-10 of training. Bottom: Trial averaged plots of speed (cm/s) aligned to cue entry on T10 for GTACR inhibition mice, separated by reward or no reward cues. Quantification of mean speed during cues. N=8 mice for mCherry, 4 for GTACR late inhibition. *p<0.05, paired t-test. N=4 mice for GTACR late inhibition. *p<0.05, paired t-test.
