## Supplemental Figure 3 for "Neural activity ramps in frontal cortex signal extended motivation during learning"

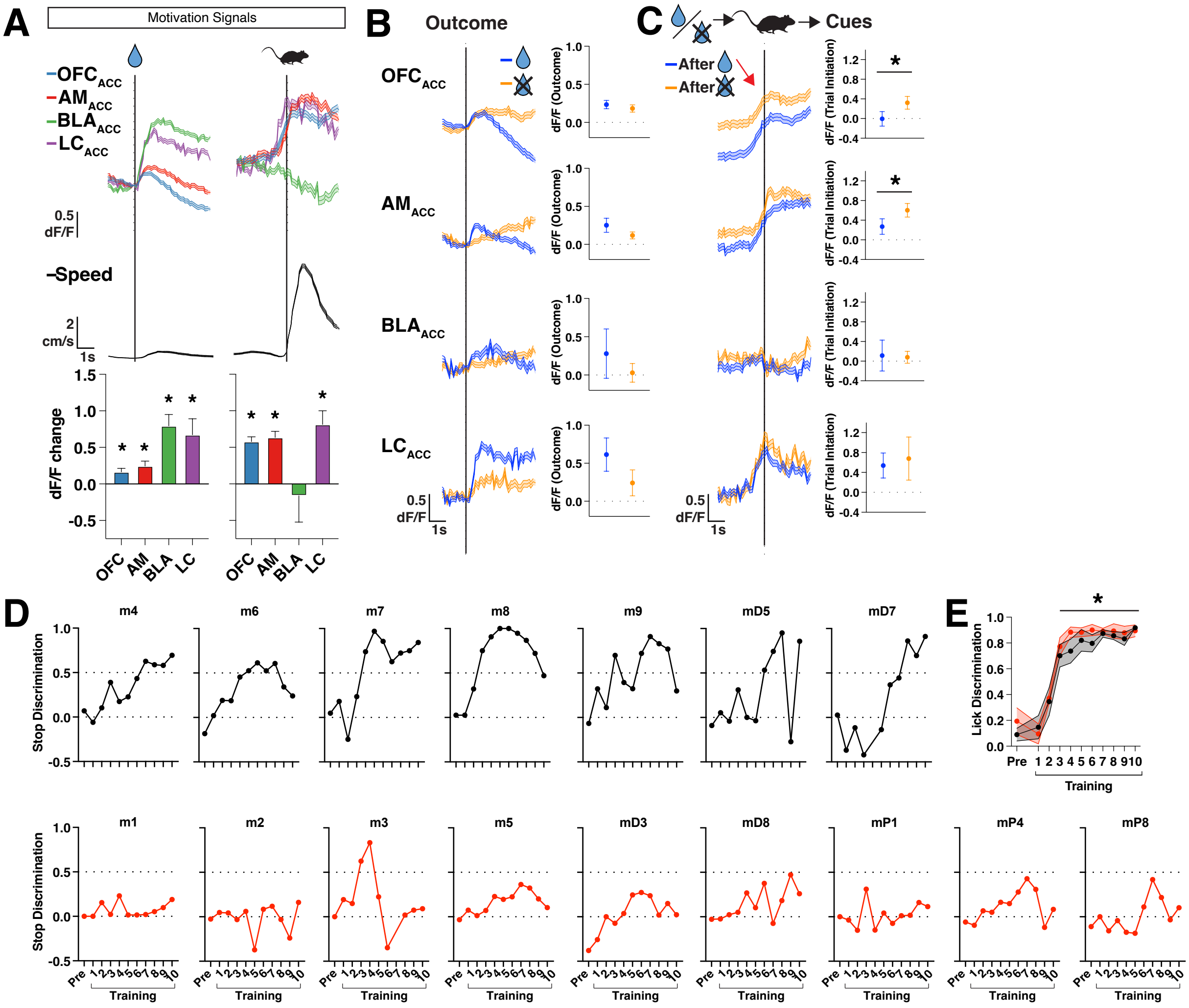


**Figure S3. Motivation signals in bulk projection activity, and behavior of learners**

(A) Top: Trial average plots of each projection activity (z-scored dF/F) and speed (cm/s) aligned to reward onset (left) or trial initiations (right). Data are mean (solid line) ± s.e.m (shaded area). Bottom: Quantification of the mean dF/F change during reward (left) or trial initiation (right; N=19, 12, 5, 4 mice). Data are mean ± SEM.

(B) Left: Trial average plots of projection activity (z-scored dF/F) aligned to outcome onset, separated by reward or no-rewards. Data are mean (solid line) ± s.e.m (shaded area). Right: Quantification of the mean dF/F during outcome. N=19, 12, 5, 4 mice, data are mean ± s.e.m.

(C) Same as C, but for trial initiations after each outcome. *$p<0.05$, paired t-test between reward and no-reward each day.

(D) Plots of stop discrimination across preexposure or training for each mouse in the “Learner” group (top) or “Non-Learner” group (bottom). Dashed line is shown for 0 and 0.5 stop discrimination.

(E) Lick discrimination between the “Learner” or “Non-Learner” mice. *$p<0.05$, one-way ANOVA between training days and preexposure, with post-doc Tukey’s multiple comparison test.
