## Supplemental Figure 4 for "Neural activity ramps in frontal cortex signal extended motivation during learning"

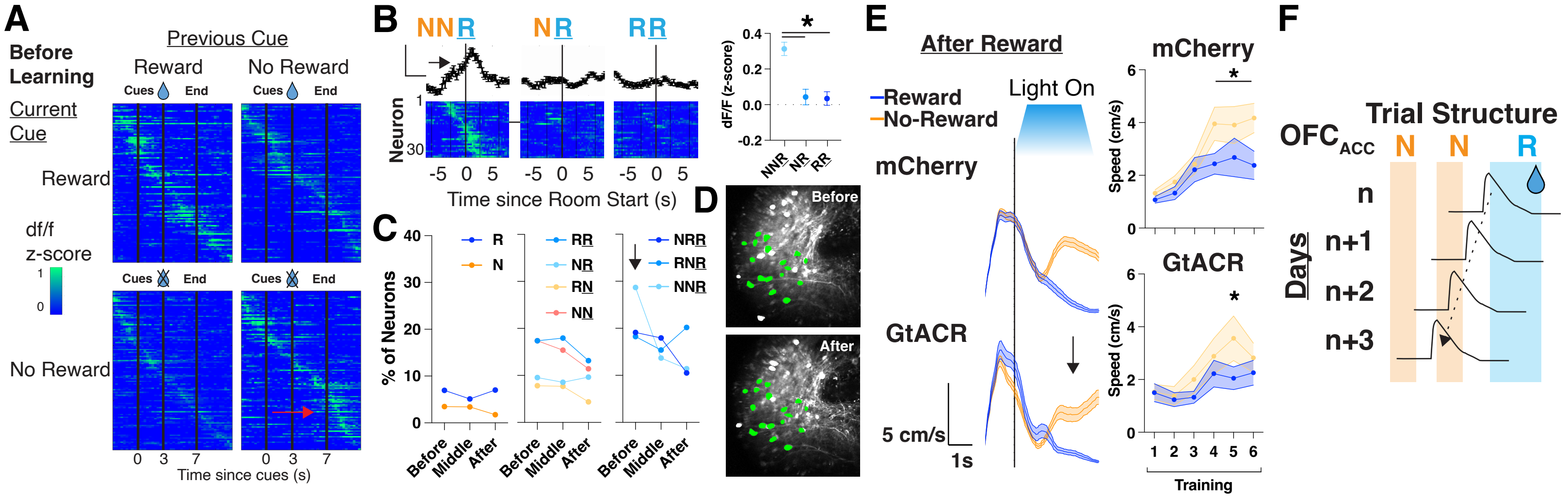


**Figure S4. Neuron tunings to NNR task structure and inhibition of $OFC_{ACC}$ neurons.**
(A) Mean z-scored dF/F for every recorded cell, aligned to cue onset separated by current cue and previous cue presentation (N=115 cells across 5 mice). Red arrow denotes the increased response after no-reward cues when the previous cue was no-reward.
(B) Left: mean population activity (top; z-scored dF/F) and individual neuron trial averaged activity (bottom) for NNR neurons during R cues preceded by 2 N, 1 N, or 1R cue presentations. Right, quantification of NNR tuned neurons' response to R cues preceded by 2 N, 1 N, or 1 R cue presentations (N=32 cells). *p<0.05, one-way repeated measures ANOVA with post-hoc Tukey's multiple comparison test.
(C) Percentage of neurons tuned (std > 0.75 3 seconds before or after cue onset) to R or N cues in total (left) or based on what cues preceded it one trial before (middle) or across two trials before (right). N=115, 117, 114 cells for "before", "middle", and "after" days. Black arrow denotes the higher percentage of neurons that are tuned to R cues after 2 N cues (NNR cells) before learning.
(D) Neural sources for a single field of view. Sources captured throughout learning are highlighted in green.
(E) Left: mean animal speed (cm/s) aligned to cue zone entry after reward on T6 for mCherry control or GtACR mice. Black arrow signifies speed increase during N cues. Right: quantification of mean change speed in cue zone after reward, assessed separately for each cue presentation. N=10 mice for mCherry and 13 mice for GtACR, *p<0.05, paired t-test.
(F) Schematic of reward-responsive OFC projection neurons becoming increasingly active during no reward cues that precede reward cues over days.
