## Supplemental Movie Caption for "Neural activity ramps in frontal cortex signal extended motivation during learning"

**Supplementary Video 1**

Behavior during learning. Playback speed: 2x. Shown here is a representative mouse learning to stop in cues that predict reward (blue walls) and run throughout consecutive cue presentations that predict no-reward (yellow walls). Displayed trial sequence order is reward, no-reward, no-reward, and reward (RNNR).
